# The Extended Life Cycle: A Multiscale Modeling Framework for Extended Evolutionary Dynamics

**DOI:** 10.1101/2025.05.18.652243

**Authors:** Nayely Vélez-Cruz, Manfred D. Laubichler

## Abstract

A complete explanation of evolutionary change requires reconciling processes that operate across multiple time scales. Development, the processes by which traits are generated, unfolds over an individual’s lifetime; heredity, encompassing the diverse forms of information transmission, occurs across generations; and population and ecological change often take place over even longer temporal horizons. Capturing these layered dynamics in a formal mathematical framework is essential for advancing evolutionary theory in line with recent conceptual developments, such as Extended Evolutionary Theory (EET) or the Extended Evolutionary Synthesis (EES). In this work, we introduce the Extended Life Cycle (ELC), a multiscale mathematical modeling framework that centers the life cycle as the fundamental unit of evolutionary analysis. In contrast to traditional gene-centric approaches, the ELC formalism captures the nested temporal structure and multilevel causal architecture of biological systems by modeling development, heredity, population, and ecological dynamics across distinct but interacting timescales. Each layer is expressed as a stochastic state-space model, enabling feedback across levels and accommodating diverse interaction topologies and inheritance systems. We demonstrate the utility of the ELC through a Bayesian simulation to estimate latent dynamics across three time scales. By formalizing the life cycle as a dynamic scaffold, the ELC offers a flexible foundation for modeling evolution in complex systems.

## 1 Introduction

In his 1956 work “Time in Biology,” J.B.S. Haldane aptly observed that many difficulties in biology arise from the need to think in terms of multiple time scales [1]. This is particularly the case for evolution, as evolutionary change is best understood through the intersection of development, heredity, and population dynamics—the key mechanisms by which traits are formed, passed down, and distributed within populations. These processes and their associated subprocesses unfold across molecular, cellular, organismal, and ecological scales, exhibiting nested structures and reciprocal feedback mechanisms [2, 3, 4, 5, 6]. Central to this multiscale perspective is the life cycle as the fundamental biological unit of analysis [7, 8] as it integrates the short-term developmental dynamics of individual organisms with long-term hereditary, population dynamic, and ecological processes. It is through the life cycle that organisms engage in various processes such as growth and development, reproduction, and adaptation and ensure the continuity of the species by linking successive generations in a population through heredity. Evolution is thus a multiscale phenomenon that emerges from the integration of development, heredity, and population dynamics, where the life cycle is the nexus of interaction between these processes.

Developing mathematical modeling tools and frameworks to capture these layered dynamics is of central interest, but not a straightforward task. From a conceptual standpoint, a challenge lies in reconciling underlying ontologies and mapping across different scales of biological organization [9]. This challenge has driven a growing interest in multiscale modeling. While multiscale modeling has become increasingly popular in a variety of biological domains, including neuroscience [10], biomedicine [11], and ecology [12], its application to the integration of development, heredity, and population dynamics may be of particular value in endowing evolutionary theory with a mathematical structure suited to match the ambitions of the more recent extended evolutionary syntheses (EESs) and extended evolution theory (EET) [6, 13, 14]. Although earlier modeling efforts have emphasized the role of developmental processes, such as developmental memory and social development, in influencing evolutionary dynamics [15, 16], more recent work has increasingly recognized the need to explicitly incorporate layered dynamics to better understand how development shapes evolutionary trajectories [17, 18]. By capturing the interdependencies across different scales, from gene regulatory network dynamics driving development to population-level changes driven by heredity and selection, multiscale modeling can provide mechanistic insights into how such processes interact to shape evolutionary outcomes. Furthermore, the growing availability of time series data across biological systems and scales, combined with methods for inferring unknown dynamical structures [19, 20, 21, 22, 23, 24, 25] and growing efforts to bridge data-driven machine learning with mechanistic multiscale modeling [26, 27, 28], enhances our ability to operationalize the life cycle through modeling frameworks that integrate development, heredity, and population dynamics.

In light of these considerations, we present a multiscale state-space modeling framework designed to integrate these three dimensions across multiple biological time scales. By leveraging state-space modeling, we capture the fine time scale dynamics occurring within an individual’s life cycle, as well as the coarse time scale dynamics that unfold across generations while accounting for reciprocal feedback between these levels. This feedback enables us to explicitly model how developmental processes shape the transmission of hereditary information across generations in a population, how development is shaped in turn by various inheritance systems, including ecological inheritance, and 3) how the environment is shaped by the actions of organisms during their life cycles. By embedding these multilevel and reciprocal causal relationships within a formal mathematical structure, our approach addresses longstanding conceptual and methodological critiques of traditional evolutionary models that treat causation as unidirectional [29] and contributes to ongoing efforts to incorporate non-genetic inheritance mechanisms into the mathematics of evolutionary theory [30]. Notably, our framework, termed the Extended Life Cycle (ELC), explicitly incorporates social development and interacting population structures without relying on traditional population models, though such models can be integrated when appropriate. It also enables the incorporation of existing mechanistic models of development [31, 32, 33, 34], heredity, and population dynamics [35], while remaining amenable to new data-driven models.

### 1.1 Development, Heredity, and Population Dynamics

Development, heredity, and population dynamics are interconnected biological processes that unfold over distinct yet deeply intertwined time scales. Development operates on the time scale of the individual organism, during which genomic, cellular, and environmental interactions, such as gene expression, hormonal signaling, and external inputs, shape phenotypic outcomes over the course of the life cycle. Moreover, development provides insight not only into the mechanisms underlying phenotypic variation and novelty [36], but also into how it structures the offspring–parent phenotype distribution [37]. These developmental processes are passed down across generations through heredity, which encompasses the diverse forms of information transmission [38, 39, 40]. Heredity not only transmits changes in development, such as shifts in timing, pathways, or rates, but this relationship is reciprocal. Development can also influence which hereditary channels (genetic, epigenetic, behavioral, or environmental) are utilized during the intergenerational transition, as seen in processes like phenotypic plasticity, genetic assimilation, and niche construction [41, 42, 43, 44, 45, 46, 47, 48]. These examples illustrate the interconnectedness of heredity and development: heritable variations in biological processes can influence developmental outcomes, while development shapes how hereditary information is expressed and transmitted. Such reciprocal causation challenges traditional gene-centric frameworks that treat inheritance as a one-way process.

Meanwhile, population dynamics occur over longer periods, where processes such as migration, drift, and natural selection describe the dynamics of variation and its evolutionary consequences on the structure of populations. Population structures establish networks of interactions that determine how hereditary channels connect, influencing which traits are transmitted across generations. These interactions can include indirect genetic effects, where the phenotype of one individual alters the phenotype and evolutionary trajectory of another [49]. For instance, the arrangement of individuals within a population can lead to varying patterns of resource sharing, social interactions, and behavioral adaptations, all of which impact how and which traits are expressed and passed on. Changes in environmental conditions or social dynamics can reshape these networks, altering the selective pressures and hereditary channels that guide trait transmission. Consequently, the connectivity within these networks plays a crucial role in influencing the pathways through which developmental processes and outcomes change or remain stable across generations.

## 2 The Extended Life Cycle

### 2.1 The ELC as the Unit of Evolution

Integrating development, heredity, and population dynamics requires a unifying concept that bridges these three dimensions, an imperative recognized by both C.H. Waddington and John Tyler Bonner [7, 50]. Historically, the life cycle has served as a central organizing idea for this purpose. E.S. Russell argued that the organism should be viewed not as a discrete entity, but as a phase within the broader continuity of the life cycle, which he regarded as the true biological individual [51, 52, 53, 54]. Bonner [7] expanded on this view, framing the life cycle as central to biology by linking evolution to changes in life cycles over time, genetics to inheritance mechanisms between cycles, and development to transformations within a single cycle, a perspective echoed in [55]. This framing has since been reaffirmed by Griffiths and Stotz [56], who have identified the life cycle as the fundamental unit for understanding biological complexity within developmental systems theory.

This convergence of theoretical perspectives positions the life cycle not solely as a conceptual metaphor, but as a biologically grounded unit of analysis, where the organism is defined by its life cycle [8, 57, 58, 55, 59, 56]. Bonner describes the life cycle as a series of steps that occur in an organized sequence, exhibiting varying degrees of disassociability, and generally comprising three phases: growth, equilibrium, and size decrease. The life cycle connects successive generations through heredity, linking the traits and processes shaped during the life cycle with the transmission of this biological information to offspring. It not only facilitates individual continuity, but serves as the mechanism through which populations persist and evolve by linking individual-level processes, such as growth and reproduction, to population-level dynamics. Populations can therefore be understood as composed of overlapping and interacting life cycles, each integrating physiological dynamics, epigenetic mechanisms, as well as social and ecological feedback. Bonner’s definition accommodates this complexity by capturing the ordered nature of life cycle phases while remaining inclusive of the diverse factors involved in each. This framing avoids an overemphasis on development alone and instead positions the life cycle as a dynamic scaffold through which both continuity and evolutionary change are realized [55].

Building on these foundational perspectives, the Extended Life Cycle (ELC) brings explicit focus to dimensions that have long been acknowledged but often remain implicit. This framework treats the life cycle as extended in two key senses: externally and temporally. Externally, the life cycle incorporates the interactions between organisms and their environments, recognizing ecological, social, and other factors as integral to the maintenance and persistence of the organism. Temporally, it spans generations through heredity, capturing the continuous transmission of biological information. Hence, we have the *extended* life cycle as the unit of evolutionary analysis. With this established, we can define the extended life cycle as the dynamic series of processes, both intrinsic and extrinsic, that are responsible for the maintenance of the organism and the reconstruction of the life cycle in the next generation. This perspective echoes previous work which integrates internal regulatory architectures with external ecological scaffolds into an evolutionary framework, wherein developmental control is distributed across internal mechanisms and structured environmental contexts [6]. Furthermore, our definition of the ELC accommodates variability in the sequence and structure of the life cycle while explicitly integrating multiple scales of biological organization.

The choice of the extended life cycle as the unit of evolutionary analysis provides both conceptual and pragmatic advantages. From a conceptual standpoint, it encompasses the spatiotemporal organization that is fundamental to organisms. Each stage of the life cycle emerges from an interacting multitude of biological processes, both internal and external to the organism, that are responsible for its maintenance and progression through time. Operationalizing the ELC formalizes the representation of these dynamics while providing a unifying context for understanding how they interact to shape phenotypic outcomes and influence evolutionary trajectories. This operationalization also lays the groundwork for understanding evolution as changes in the life cycle over time, a perspective that naturally incorporates extended inheritance by accounting for the diverse hereditary mechanisms of information transmission responsible for introducing variation into life cycles.

Furthermore, the adoption of the ELC broadens the scope of evolutionary biology by addressing phenomena that classical gene-centric frameworks cannot fully explain. Evolutionary biology seeks to understand the types of variation that drive phenotypic change, the mechanisms by which this variation is transmitted across generations, and how phenotypic variation arises. Within the gene-centric framework, these questions are typically addressed via genetic variation, Mendelian inheritance, and genetic mutations. To take cancer evolution as an example, the ELC broadens our understanding by emphasizing the dynamic, multiscale processes that shape tumor progression. Instead of focusing solely on genetic mutations, this perspective highlights how stages in the cancer life cycle, such as cell proliferation, adaptation to the microenvironment, and metastasis, interact with environmental pressures to drive evolution. Non-genetic mechanisms, such as epigenetic modifications and phenotypic plasticity, enable cancer cells to inherit adaptive traits and respond to stressors like therapy without requiring immediate genetic changes [60, 61, 62, 63]. Moreover, cancer evolution is inherently multiscale, with selection acting on individual cells, tumor tissues, and their interactions with the host, illustrating the interconnectedness of processes across scales. This is but one example demonstrating that life cycles are the arenas where genetic, epigenetic, and ecological mechanisms interact to shape evolutionary outcomes.

Not only can the ELC offer a more comprehensive account of evolution by incorporating non-genetic forms of inheritance, ecological mechanisms, and multiscale interactions, but it also expands the scope of evolutionary theory into the realm of complexity science. As a multiscale system, the ELC facilitates the exploration of emergent properties of systems, such as robustness, evolvability, modularity, stability, and the control of variation, under the unifying umbrella of the life cycle [64, 65, 66]. Examining these complex systems’ properties as properties of the ELC enhances our understanding of evolutionary processes by demonstrating, for instance, how robustness buffers life cycles against perturbations, enabling them to persist over time, how modularity facilitates the adaptation of specific traits without destabilizing the entire system, how evolvability enables systems to generate novelty and explore adaptive possibilities, and how the control of variation balances constraint and flexibility in evolutionary trajectories.

From a practical standpoint, treating the ELC as the unit of evolutionary analysis enables the examination of specific subcomponents, such as individual traits, developmental stages, and processes (e.g. physiological changes, gene regulatory mechanisms, and developmental pathways) that collectively drive the dynamics of individual organisms while remaining integrated within the broader context of the life cycle. These subcomponents can be observed and measured through time series data collected across different biological scales. By embedding these time series within the ELC framework, researchers can model how interactions among developmental, physiological, and molecular processes shape broader evolutionary patterns. Contextualizing these dynamics within the spatiotemporal organization of the life cycle provides a cohesive foundation for integrating and interpreting such data, enabling a multiscale perspective that ultimately connects development, heredity, and population dynamics.

To fully explore the implications of the life cycle as the central unit of evolutionary analysis, a mathematical modeling framework is necessary to formalize its conceptual foundations and operationalize its application. Such a framework must therefore account for the multiscale structure of the life cycle, with processes operating on distinct but nested time scales. It should also reflect the dynamic interdependencies among these scales, incorporating feedback mechanisms that link processes across different levels of biological organization. Additionally, the modeling framework should account for the stochastic nature of biological systems, which influences their dynamics and degree of variability. It should also be capable of representing interaction networks, such as gene regulatory networks (GRNs), metabolic pathways, and population structures. Embedding these dynamics within a predictive framework enables the connection between conceptual insights of the ELC and empirical data, allowing for the integration of real-world time series into multiscale models. These requirements establish the foundation for a multiscale modeling framework capable of advancing our understanding of evolution via the ELC.

### 2.2 ELC Multiscale Modeling Framework

In light of the nested structure of development, heredity, and population dynamics, our modeling framework addresses several critical requirements: 1) a multiscale structure that captures dynamics across multiple time scales, 2) feedback within and between scales, 3) representation of interaction networks (e.g. population structures, GRNs, metabolic networks), 4) incorporation of stochasticity, and 5) the ability to integrate real-world data. Our framework employs state-space modeling [67, 68], which offers several significant advantages. Firstly, state-space models are representations of dynamical systems consisting of time-series latent states and measurements, where simulated or real-world measurements are used to infer any unknowns. Secondly, these unknowns can be inferred using Bayesian learning methods [69, 70, 71, 72, 73, 74], allowing for the development of predictive models from time-series data. Lastly, state-space models offer flexibility in their design, enabling researchers to incorporate linear or nonlinear relationships, various types of stochasticity, structures, and control inputs [75, 76, 77, 78, 79, 80].

Our modeling framework, depicted in Figure 1, captures dynamics evolving across *L* nested time scales. At each scale *l* ∈ {1, …, *L*} finer-scale time series (*l < l* + 1) are embedded within the coarser-scale dynamics. To represent multiscale feedback, each state at scale *l* depends not only on its own past but also on the coarser-scale states at *l*^′^ *> l* evaluated at a designated time point(s) 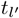, and on the full trajectories of finer scales *l*^′^ *< l* across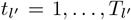. Within this setup, we consider a population of *D* individuals. This approach is akin to individual-based modeling (IBM), where each entity is explicitly modeled [81, 82, 83, 84, 85]. Such models are especially effective in capturing individual heterogeneity, local interactions, and emergent population-level patterns in ecological systems [86, 87, 88, 89]. Let 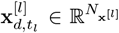 denote the latent state vector for individual *d* at time point *t*_*l*_ in time scale *l*, where each *l* consists of *t*_*l*_ = 1, …, *T*_*l*_ time points. Let 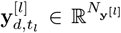 denote the corresponding measurement vector. The state-space formulation is given by Equations 1 and 2.

**Figure 1.**
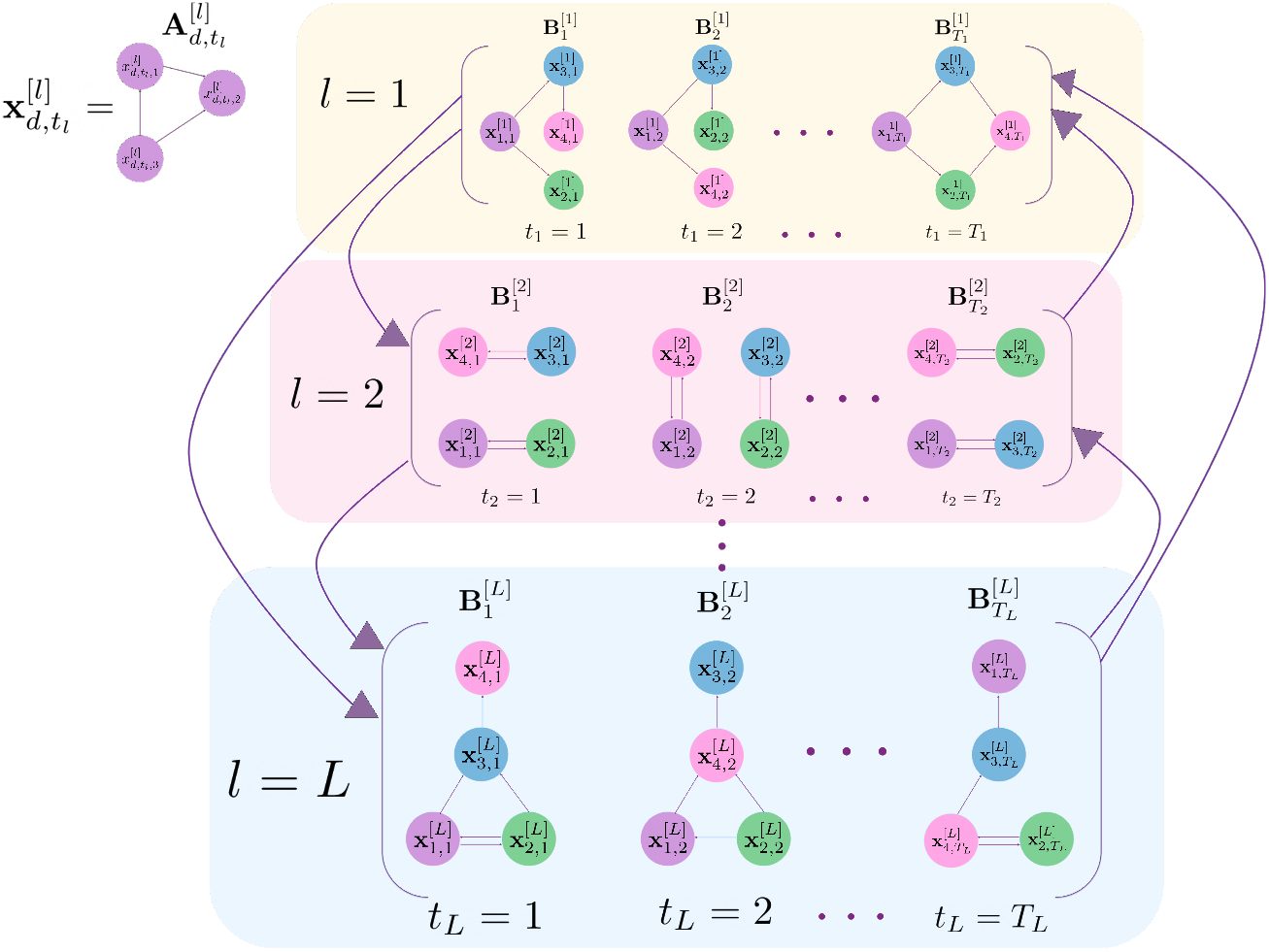
Illustration of the Extended Life Cycle (ELC) mathematical modeling framework. The diagram depicts interactions across nested time scales, highlighting both within- and cross-scale feedback. The relationships between entries in the vector 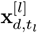 are encoded by a possibly time-varying matrix 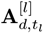. Each layer *l* depicts relationships between *d* individuals whose interactions are encoded through the possibly time-varying matrix 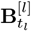.

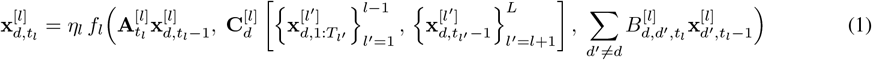

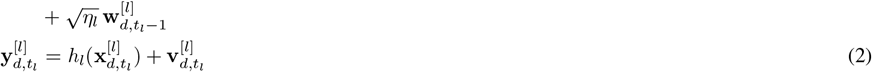

Here, *η*_*l*_ is a scale-dependent coefficient used to modulate the rate of change at each scale consistent with slow-fast systems theory [90]. The matrix 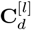 contains scale-specific coefficients that control how states at other time scales influence dynamics at scale *l*. Relationships between components in the state vector 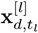 interactions are captured by the possible time-varying adjacency matrix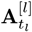, which may, for example, represent gene regulatory or pleiotropic constraints [91]. Inter-individual interactions are captured by 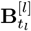 with entries 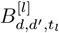, a possibly time-varying matrix encoding interactions between individuals *d* and *d*^′^ ≠ *d*, such as social behaviors, maternal provisioning, or ecological competition.

### 2.3 Modeling Development, Heredity, and Population Dynamics

To illustrate how our multiscale modeling approach can be applied to capture developmental, hereditary, and population dynamics, we consider *L* = 2 time scales: a fine time scale for individual-level life cycle changes and a coarse time scale for hereditary processes that shape population-level dynamics. More specifically, let the fine time scale, denoted by time index *t*_1_, capture the dynamics occurring within the life cycle of an organism, such as development, growth, or other within-individual-level processes of interest. Developmental processes at this scale can be seen as constructing the phenotype over time in response to genetic, epigenetic, and environmental influences, a view articulated in early conceptual frameworks such as Waddington’s notion of the “epigenetic landscape” [92, 93, 94]. Recent advances have demonstrated that incorporating explicit models of developmental dynamics can substantially improve evolutionary prediction and inference [95]. The coarse time scale time index *t*_2_ captures intergenerational change over time. Let 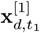 denote the life cycle state of individual *d* ∈ {1, …, *D*} at time *t*_1_ in generation *t*_2_. Let 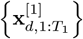 denote the developmental trajectory for all time points 1, …, *T*_1_. For example, the set of gene expression values, morphogenic growth stage, epigenetic modifications, genotypic information, metabolic activity levels, or cellular differentiation states at time *t*_1_ can be represented by 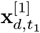. Our modeling approach aligns with the view that developmental trajectories can themselves be inherited as dynamic processes, reconstructed through regulatory interactions across generations [96]. Similarly, let 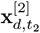 denote the hereditary state of individual *d* ∈ {1, …, *D*} at coarse time point *t*_2_, such as sets of genes, ecological niche states, maternal provisioning traits or learned skills. At the fine time scale *l* = 1, the latent state evolves according to Equation 3:

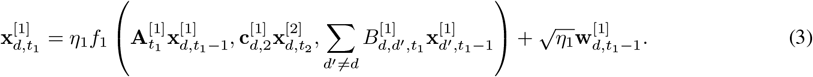

Here, 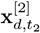 represents coarser-scale hereditary factors, such as genotypes or epigenetic states, that influence the developmental dynamics at the finer *t*_1_ time scale. The network 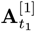 can encode a variety of relationships, including signaling pathways that affect cellular differentiation or intercellular communication during a particular stage *t*_1_ in the life cycle. The matrix 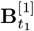 captures socially mediated developmental effects akin to extra-genetic inheritance, as in Jablonka and Lamb [13], by allowing an individual’s development to depend on the life cycle trajectories of others. More broadly, 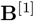 enables the modeling of indirect genetic effects (IGEs) and social selection [97], where the phenotype or genotype of one individual influences both the expression and evolutionary outcomes of traits in others. At the intergenerational (coarse) time scale *l* = 2, the latent state depends on the full trajectory of scale *l* = 1, whose dynamics are given in Equation 4.

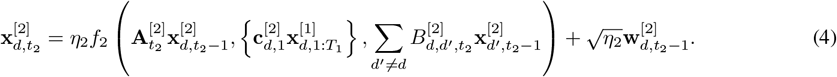

The matrix 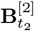, with entries 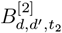, encodes relationships between individuals that influence heredity or trait transmission across generations. It may represent mating partnerships (with elements indicating probabilities or strengths of genetic exchange), kinship structures (reflecting genetic similarity), or social alliances that shape patterns of genetic exchange or trait reinforcement across a population. In contrast, the matrix 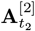 captures internal, within-lineage factors that influence how hereditary factors evolve over time. As 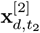 denotes hereditary factors, 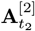 may encode genetic linkage patterns, selective pressures, or developmental constraints that affect the co-inheritance and transformation of trait values within an individual’s lineage. Lastly, unlike models that assume developmentally independent genotypes [17], our framework allows for feedback from the developmental trajectories 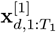 to influence hereditary factor dynamics, consistent with the principles of genetic accommodation [36]. The corresponding measurements are given by Equations 5-6.

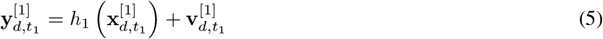

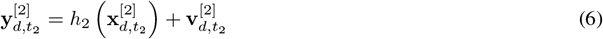

At the short-term level 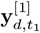, data could include time series such as body temperature, heart rate, or activity levels recorded daily or weekly. At the intergenerational level 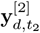, data might capture metrics like lineage-based survival rates, reproductive timing, trait persistence across generations, or genealogical records tracking phenotypic changes over time. Ecological feedback can be incorporated through addition of a third layer, which we present in Section 3.

## 3 Application to Life Cycle Evolution

Having detailed the general ELC multiscale modeling architecture, we now present a specific instantiation that demonstrates how the ELC can be utilized in practice. This application highlights how the key conceptual principles of the ELC can be operationalized to explore evolutionary dynamics across three time scales. Specifically, the first layer and finest time scale describes the life cycles of individual organisms. The second layer and intermediate time scale describes intergenerational change over time via heredity. Lastly, the third layer and coarsest time scale models niche construction dynamics, with feedback between each of the layers.

### 3.1 Individual Life Cycle Model

The model in layer *l* = 1 captures dynamics within an individual’s life cycle, modeling changes in size, reproductive effort, and survival probability. It draws conceptual inspiration from optimal control theory approaches to life history tradeoffs between survival, growth, and reproduction, now embedded within a multiscale framework [98, 99, 100]. The general formulation is given in Equation 7:

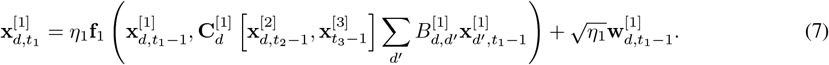

The state vector 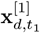 comprises size 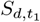, reproductive effort 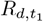, and survival probability 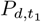 (Equation 8):

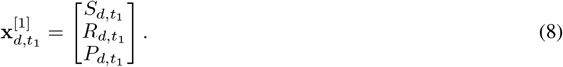

Here, **f**_1_ is a vector-valued function defining the dynamics of 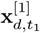. Developmental influences are modeled as nonlinear mappings from hereditary factors 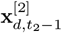 and environmental conditions 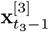, consistent with growing emphasis in quantitative genetics on nonlinear genotype–phenotype maps [101]. Hereditary inputs include factors influencing size 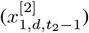, reproductive effort 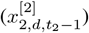, and survival 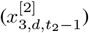, while environmental influences may include nutrient availability or resource levels [102, 103, 104, 105, 106]. Interactions between individuals are encoded by 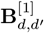, and stochastic variation is captured by the noise term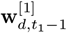.

Growth is influenced by genetic and developmental constraints, resource availability, and competitive interactions, with additional modulation by age structure, environmental variation, and stochastic effects [107, 108]. To model these dynamics, we define a growth equation for size 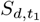 as a function of hereditary factor 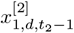, environmental conditions 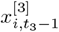 in niche *i*, and competition via the interaction matrix **B**^[1]^:

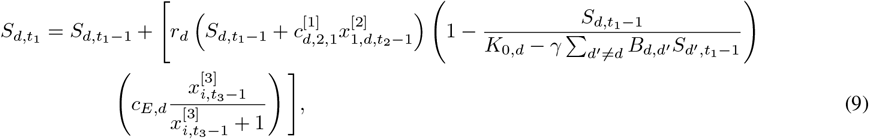

where *r*_*d*_ is the intrinsic growth rate, 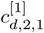 and *c*_*E,d*_ scale hereditary and environmental influences, respectively, and *γ* modulates competitive pressure via 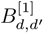. The environmental term follows a Michaelis–Menten-type saturation function, widely used to model diminishing returns in ecological and physiological systems [109]. The individual’s maximum size *K*_0,*d*_ is modulated by competitive effects from neighbors, with interaction intensity governed by *γ* and the entries in **B**^[1]^.

Reproductive effort 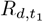 is influenced by body size, hereditary factors, environmental influences, and tradeoffs between survival and reproduction. The model is given by Equation 10.

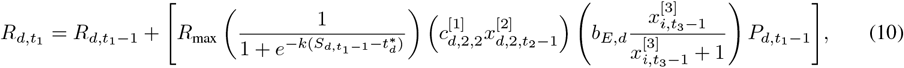

where *R*_max_ denotes the maximum potential reproductive investment, 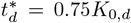 is the threshold size above which reproductive investment increases sharply, and *k* controls the steepness of this sigmoidal transition. The term 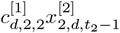 reflects the effect of intrinsic hereditary factors (such as fecundity or reproductive timing), where 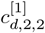 scales the hereditary factor effect. The term 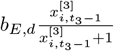 captures the saturating effect of environmental resources. Reproductive effort is also scaled by the current survival probability 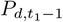, so that higher 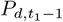 increases reproductive effort. In this formulation, individuals delay significant reproductive investment until they reach a species- or strategy-specific threshold size, consistent with the idea that reproduction is energetically costly and requires sufficient somatic growth [110]. This also captures the idea that life history strategies depend on state variables that incorporate past energy allocation decisions through dependence on 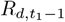[111].

Survival probability 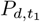 is influenced by size, environment, and reproductive effort. Rather than using a classical hazard function, we define a protection score 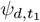 that captures survival-enhancing and survival-reducing influences:

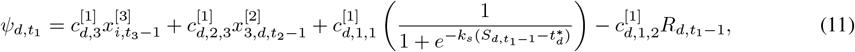

where scaling factors 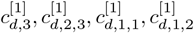 modulate environmental effects, survival-related traits, size contribution, and reproductive costs. The size effect follows a sigmoidal curve centered at 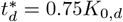, capturing a threshold-like transition: limited protection below this point, then rapidly increasing survival benefit before saturation. Survival is then modeled via Equation 12.

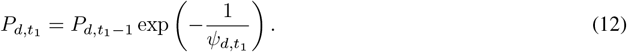

This form captures the trade-off between growth, reproduction, and survival: larger size and better environment increase protection, while reproduction incurs survival costs.

#### 3.1.1 Intergenerational Dynamics

In layer *l* = 2, we model the intergenerational transmission of traits over time scale *t*_2_ using a modified Hansen model [112, 113, 114, 115]. It models change in quantitative traits as an Ornstein Uhlenbeck process and has been used to model genetic drift with stabilizing selection as well as adaptive evolution with shifting optima [116]. The general formulation of the model is given by Equation 13.

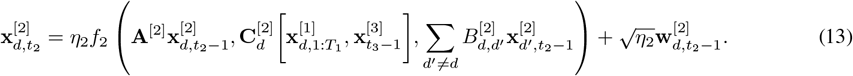

In this setup, the matrix **A**^[2]^ encodes linkage relationships between components in the hereditary state 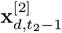. In this layer, *d* can be thought of as a lineage. Note that 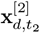 is also a function of the individual life cycle trajectory, the state of the environment 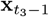, and the traits of individuals *d*^′^ with whom *d* interacts with through the matrix **B**^[2]^. Consistent with perspectives emphasizing that developmental processes and environmental factors play active, constructive roles in shaping evolution [117], these dependencies allow hereditary change to be dynamically shaped by both internal and external developmental contexts. Lastly, 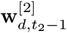 is a noise vector. The specific functional form of *f*_2_ is given by Equation 14.

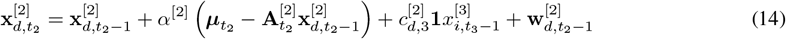

where *α*^[2]^ is the update rate controlling the strength of reversion to the trait target ***µ***, and the term 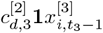 introduces an environmental influence term applied to all trait dimensions. Here, **1** is a vector of ones matching the dimensionality of the trait vector **x**^[2]^.

To model adaptive evolution, the dynamic target 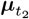 is computed in Equation 15 from a baseline trait value ***µ***_*d*,0_ and two additional components: a weighted summary of the individual’s life cycle trajectory and an interaction term capturing the influence of other individuals’ life cycle trajectories. This captures the idea that plasticity can influence not only phenotypic outcomes but also the rate and direction of evolutionary change [118].

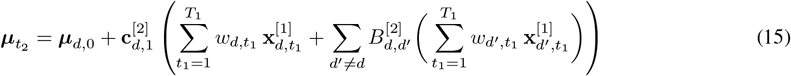

This formulation allows 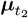 to dynamically respond both to the individual’s life history and to structured interactions with other individuals, as may occur through kin, social, or ecological feedback mechanisms.

#### 3.1.2 Layer 3: Niche Construction Dynamics

Layer *l* = 3 models the dynamics of the environmental state vector 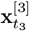, which reflects ecological components such as resource levels or temperature. These dynamics are shaped by feedback from individual-level life cycles (Layer 1) and hereditary factors (Layer 2), each scaled by 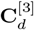, and are updated according to:

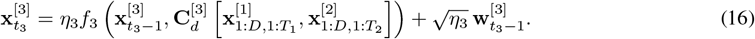

The function *f*_3_ combines baseline recovery, density-dependent regeneration, and feedback from individual contributions and consumption:

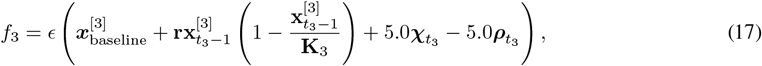

where **r** is the regeneration rate, **K**_3_ the carrying capacity, and *ϵ* the update rate on timescale *t*_3_. The logistic term models resource renewal slowing near saturation. The contribution 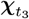 and consumption 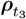 vectors are computed by aggregating individual-level activity across assigned environmental components. For each individual *d*, we first compute a combined average from life cycle and hereditary state variables *x*^[1]^ and hereditary traits *x*^[2]^ at time *t*_3_:

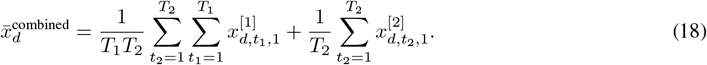

The population mean at *t*_3_ is then given by:

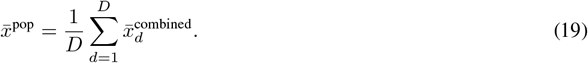

The deviation from the population mean is defined as:

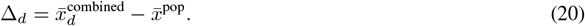

We then compute the individual’s contribution and consumption scores as:

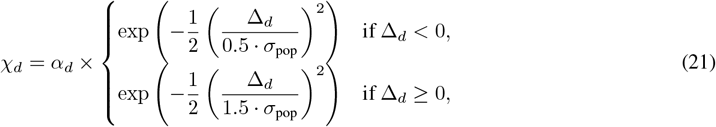

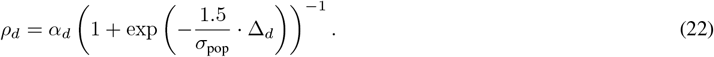

Here, *σ*_pop_ is the standard deviation of 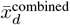 across the population at *t*_3_, used to tune the sharpness of the Gaussian and sigmoid functions. Individuals below the mean are penalized more steeply in contribution score, while consumption saturates for individuals with large positive deviations. Both scores are scaled by *α*_*d*_, an individual-specific weight. Total scores for each environmental component *i* are obtained by summing over individuals assigned to it:

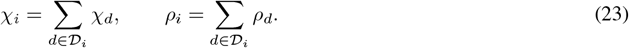

The values *χ*_*i*_ and *ρ*_*i*_ are then used in Equation 17 to compute environmental change. This formulation captures how individual developmental and hereditary factors collectively result in niche construction dynamics.

## 4 Simulations

Simulations provide a tractable and controlled setting to illustrate the ELC framework. We demonstrate how the ELC can be utilized in practice using a simple Bayesian application. While the scenarios presented here represent only a small subset of its potential applications, they demonstrate the framework’s capacity to capture multiscale evolutionary dynamics. We simulate noisy measurements at each time scale and estimate the latent state trajectories using a multiscale particle filtering approach. Rather than testing a specific hypothesis, this simulation demonstrates how the model captures conditions under which emergent evolutionary patterns can arise. The simulations illustrate several emergent patterns resulting from multiscale feedback: (1) size, survival, and reproductive outcomes shaped by the nonlinear integration of developmental, hereditary, and ecological influences, (2) dynamic divergence and convergence of traits driven by developmental, hereditary, and environmental feedback, and (3) life cycle-mediated niche construction.

We consider a small community consisting of *D* = 4 individuals in varying environments whose life cycle dynamics evolve according to Equations (7, 9-12) over the *t*_1_ time scale, which captures growth 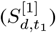, reproductive effort 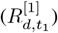, and survival probability 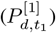, where 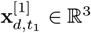 is the state vector defined according to Equation (8). Over the intergenerational time scale *t*_2_, the dynamics consider trait evolution in 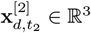 across generations and are given by Equations (13-15). Lastly, niche construction dynamics are described by the evolution of the environmental state 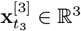 in the coarsest time scale, *t*_3_, and are governed by Equations (16-23). For each simulation, we set *T*_1_ = 51, *T*_2_ = 21, and *T*_3_ = 11 time points where all *T*_1_ = 1, …, 21 are nested in each *t*_2_, and all *T*_2_ = 1, …, 21 are nested in each *t*_3_. The code is publicly available on GitHub at https://github.com/nvelezcruz/Extended-Life-Cycle-PF.

## 5 Simulation Example

In this scenario, at time *t*_3_ = 0, each individual *d* is placed into a favorable or a harsh environment, as is specified by the baseline environmental state 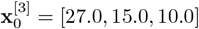. Individual *d* = 1 is assigned to component 1, individual *d* = 3 is assigned to component 2, and individuals *d* = 2 and *d* = 4 are assigned to component 3. At *t*_3_ = *T*_3_*/*2, this baseline switches from 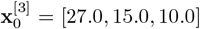 to 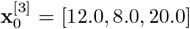 so that individuals *d* = 1 and *d* = 3, who were previously in favorable environments are placed into harsh environments (components 2 and 3, respectively), while individuals *d* = 2 and *d* = 4, who were previously in harsh environments, move to a favorable environment (component 1). This simulation captures phenomena such as environmental fluctuations or habitat changes, where individuals experience dynamic shifts in their surroundings. For the finest time scale *t*_1_, each individual’s state is initialized with specific values for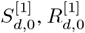, and 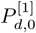, as shown in Table 1, alongside the parameters governing their dynamics. Individuals in favorable environments (individuals *d* = 1 and *d* = 3) have higher initial sizes 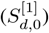 and growth rates 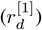, and greater initial maximum sizes 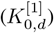 than individuals in harsh environments (Individuals 2 and 4). Reproductive efficiency 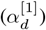, environmental influence coefficients 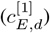, and trade-offs between reproduction and survival 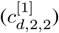 are also higher for individuals in favorable environments and lower for those in harsh environments. The reproductive effort parameters are given in Table 2. Next, the interaction matrix 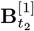 captures direct effects between individuals during their life cycle. We restrict this matrix so that only individuals in the same type of environment can interact (harsh versus favorable) despite potentially being in different components of 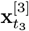. For simplicity, we remove the temporal evolution. It is defined as:

**Table 1:** Layer 1: Initial conditions and individual-specific parameters used in the simulation.

| Ind. | Env. Comp. | $S_{d,0}^{[1]}$ | $R_{d,0}^{[1]}$ | $P_{d,0}^{[1]}$ | $r_d$ | $K_{0,d}$ | $c_{E,d}^{[1]}$ | $\alpha_d^{[1]}$ | $c_{d,3}^{[1]}$ | $c_{d,2,3}^{[1]}$ | $c_{d,1,1}^{[1]}$ | $c_{d,1,2}^{[1]}$ |
| --- | --- | --- | --- | --- | --- | --- | --- | --- | --- | --- | --- | --- |
| 0 | 1 (Favorable) | 10 | 0.1 | 1.0 | 0.07 | 40 | 0.6 | 1.0 | 1.0 | 1.0 | 1.0 | 20.0 |
| 1 | 3 (Harsh) | 5 | 0.1 | 1.0 | 0.04 | 20 | 0.5 | 1.0 | 1.0 | 1.0 | 1.0 | 20.0 |
| 2 | 2 (Favorable) | 10 | 0.1 | 1.0 | 0.06 | 30 | 0.6 | 1.0 | 1.0 | 1.0 | 1.0 | 20.0 |
| 3 | 3 (Harsh) | 5 | 0.1 | 1.0 | 0.03 | 15 | 0.5 | 1.0 | 1.0 | 1.0 | 1.0 | 20.0 |

**Table 2:**
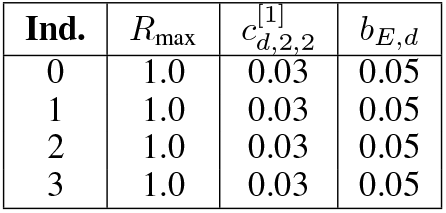
Reproductive effort parameters for each individual.

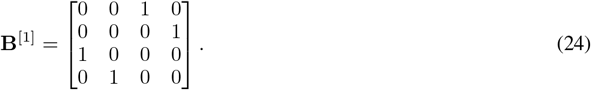

Lastly, the process noise is modeled as additive Gaussian noise **w**^[1]^ ∼ 𝒩 (**0**, (0.01)^2^**I**_3_).

For Layer 2, hereditary factors 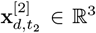 represent latent traits that influence individual life cycle dynamics, including growth, reproductive effort, and survival. These traits are initialized using individual-specific baselines 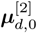 (Table 3) and evolve over time according to a modified Ornstein–Uhlenbeck process as defined in Equations (13)–(15). Trait dynamics incorporate both individual life history feedback (from Layer 1) and trait-level interactions across individuals via an interaction matrix **B**^[2]^. The trait update equation is governed by the reversion rate *α*^[2]^ = 0.5, with the dynamic target 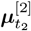 incorporating weighted developmental summaries and interaction effects. Environmental modulation is introduced through an additive term 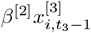, where the environmental effect is scaled by an individual-specific coefficient 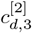. Gaussian noise 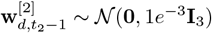 is added at each step to reflect stochastic variation. Weights 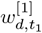 and 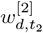 used in the computation of 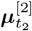 are generated by sampling from a uniform distribution and normalized such that they sum to 1 across each time axis. The hereditary interaction matrix **B**^[2]^ mirrors the life cycle interaction matrix **B**^[1]^, encoding competitive interactions between individuals occupying similar environments. Internal trait interactions are governed by the matrix **A**^[2]^:

**Table 3:**
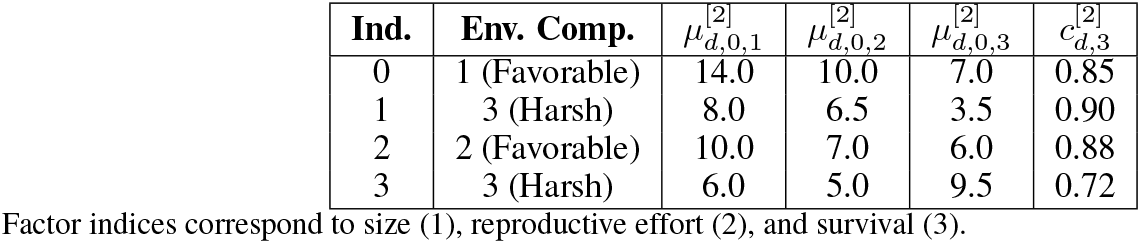
Layer 2 initial hereditary factor baselines and parameters. Global parameters are listed per individual for clarity.

| Ind. | Env. Comp. | $\mu_{d,0,1}^{[2]}$ | $\mu_{d,0,2}^{[2]}$ | $\mu_{d,0,3}^{[2]}$ | $c_{d,3}^{[2]}$ |
| --- | --- | --- | --- | --- | --- |
| 0 | 1 (Favorable) | 14.0 | 10.0 | 7.0 | 0.85 |
| 1 | 3 (Harsh) | 8.0 | 6.5 | 3.5 | 0.90 |
| 2 | 2 (Favorable) | 10.0 | 7.0 | 6.0 | 0.88 |
| 3 | 3 (Harsh) | 6.0 | 5.0 | 9.5 | 0.72 |

Factor indices correspond to size (1), reproductive effort (2), and survival (3).

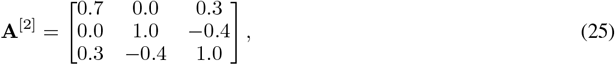

where each row corresponds to one of the hereditary factors—size, reproductive effort, and survival—and each column represents the influence of a given factor on the factor in that row. This matrix defines how an individual’s factors regulate one another during heredity. In the first row, the size trait is positively influenced by its own previous value (0.7), reflecting strong trait-level persistence, and by the survival trait (0.3), indicating that greater baseline survivability may contribute to somatic growth or maintenance. There is no direct influence from reproductive effort, as indicated by the zero entry. In the second row, reproductive effort is entirely self-regulated (1.0), and negatively influenced by survival (−0.4). This implies that individuals with higher baseline survival may reduce their reproductive allocation, consistent with life-history strategies in which long-lived individuals reproduce more conservatively. The size trait does not directly influence reproductive effort. The third row shows that survival is positively influenced by body size (0.3), negatively influenced by reproductive effort (−0.4), and also exhibits strong self-regulation (1.0). This structure reflects a classical trade-off, where larger size may promote survival through increased robustness or resource access, but higher reproductive effort reduces survival due to energetic and physiological costs.

Niche construction dynamics at the *t*_3_ scale are governed by the environmental state vector 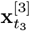, which tracks the values of three abiotic or resource components. At *t*_3_ = 0, the environment is initialized as 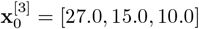]. At *t*_3_ = *T*_3_*/*2, an abrupt shift to [12.0, 8.0, 30.0] is introduced, modeling a regime change that redistributes environmental conditions across components. The regeneration of each component follows logistic growth, with carrying capacities **K**_3_ = [25.0, 18.0, 12.0] and regeneration rates **r**_*x*_ = [0.4, 0.8, 0.6], capturing baseline recovery dynamics in the absence of feedback. Environmental updates are shaped by both individual-level contributions 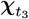 and consumptions 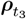, which are computed from deviations in average trait expression relative to the population. For these simulations, each individual’s deviation is calculated from a combined average of life cycle states corresponding to size 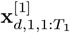 and hereditary traits 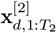 as an optional parameter. The deviations are scaled by population variance to produce asymmetric Gaussian and sigmoid-based scores. In larger-scale scenarios with many individuals, the population mean used to compute these deviations would typically be calculated using only those individuals assigned to the same environmental component. However, since we are considering a small population in this simulation, we compute the mean using all individuals across all components. To isolate the effects of life cycle dynamics, hereditary contributions 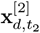 are set to zero when computing trait averages in this simulation; this is a modeling choice specific to the current analysis and can be adjusted in the code. Individuals are assigned to environmental components as follows: component 1 receives contributions from individual *d* = 1, component 2 from *d* = 3, and component 3 from *d* = 2 and *d* = 4. Individual impact on the environment is scaled by weights 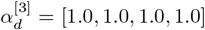. Time-averaged trait values are computed across *t*_1_ and *t*_2_ using normalized random weights 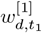 and 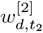. Contributions and consumptions are aggregated per component and passed into the environmental dynamics model to determine the next state 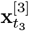, as described in Equation (17).

### 5.1 Particle Filter Parameters

To demonstrate how the multiscale models developed above can be used within a Bayesian inference framework, we assume that the latent state trajectories 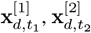, and 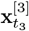 are unknown and must be inferred from noisy measurements. We use a particle filtering approach (Algorithm 1) to estimate these hidden states across all time scales. The particle filter is run with *N*_particles_ = 50 particles. At each layer, we assume additive Gaussian measurement noise. The standard deviations for the observation models are

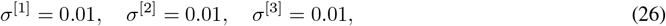

corresponding to the observations 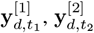, and 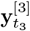 respectively. To simulate the measurements, vector-valued Gaussian noise is added to each latent state:

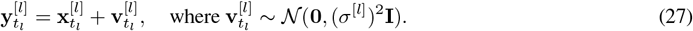

We also assume additive Gaussian process noise at each layer, with covariance given by 1*e*^−3^**I**_3_, to reflect uncertainty in the latent dynamics:

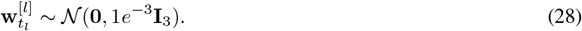

This setup enables us to evaluate the particle filter’s performance in recovering multiscale trajectories under known noise conditions.

#### Algorithm 1

Multiscale Particle Filter

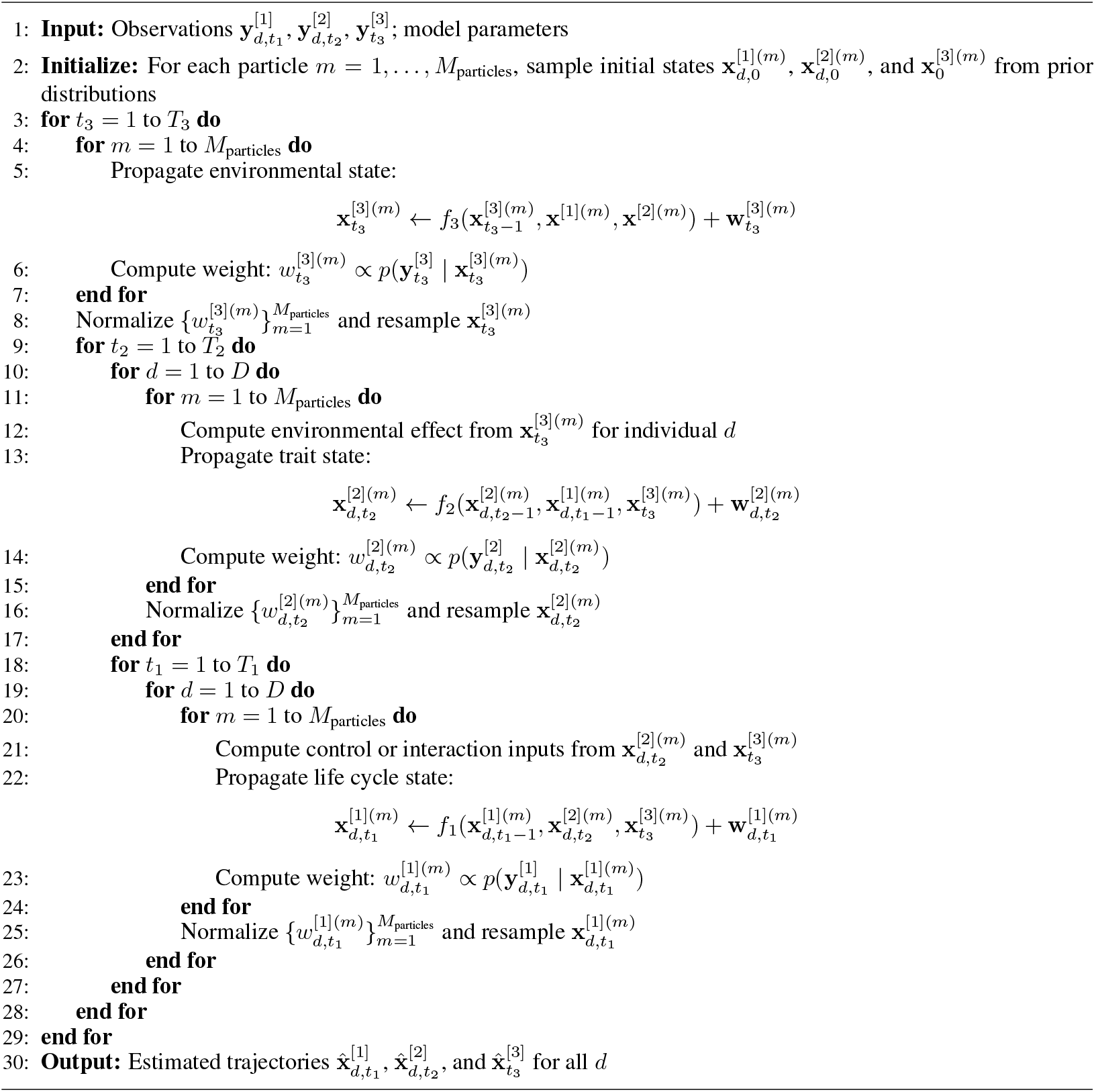

### 5.2 Results and Discussion

#### 5.2.1 Layer 1: Individual Life Cycle Dynamics

Figures 2(a) and 2(b) depict the true versus estimated size dynamics for each individual *d*. Specifically, Figure 2(a) shows the size dynamics prior to the environmental shift at *t*_3_ = 1 and *t*_2_ = 20, whereas Figure 2(b) shows the size dynamics following the environmental shift at *t*_3_ = 10 and *t*_2_ = 20. Due to competitive interactions, neither individual *d* = 1 nor *d* = 3 is able to reach their respective maximum body size values (40 and 30, respectively). Similarly, individuals *d* = 2 and *d* = 4 fall below their respective maximum sizes (20 and 15, respectively), reflecting the influence of competition and resource limitation. A comparison of Figures 2(a) and 2(b) further reveals that the rate of growth for individuals *d* = 1 and *d* = 3, who transitioned from resource-abundant to resource-deficient environments, exhibits a small decline following the environmental shift. Conversely, individuals *d* = 2 and *d* = 4, who moved from resource-deficient to resource-abundant environments, show a slight increase in their growth rates after the shift. The influence of the hereditary factor 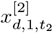 contributes additively to growth, helping to sustain growth trajectories even under shifting environmental conditions. This is akin to an additive genetic effect in quantitative genetics moderating the impact of environmental fluctuations on individual size dynamics, impacting growth in both resource-rich and resource-poor settings.

**Figure 2.**
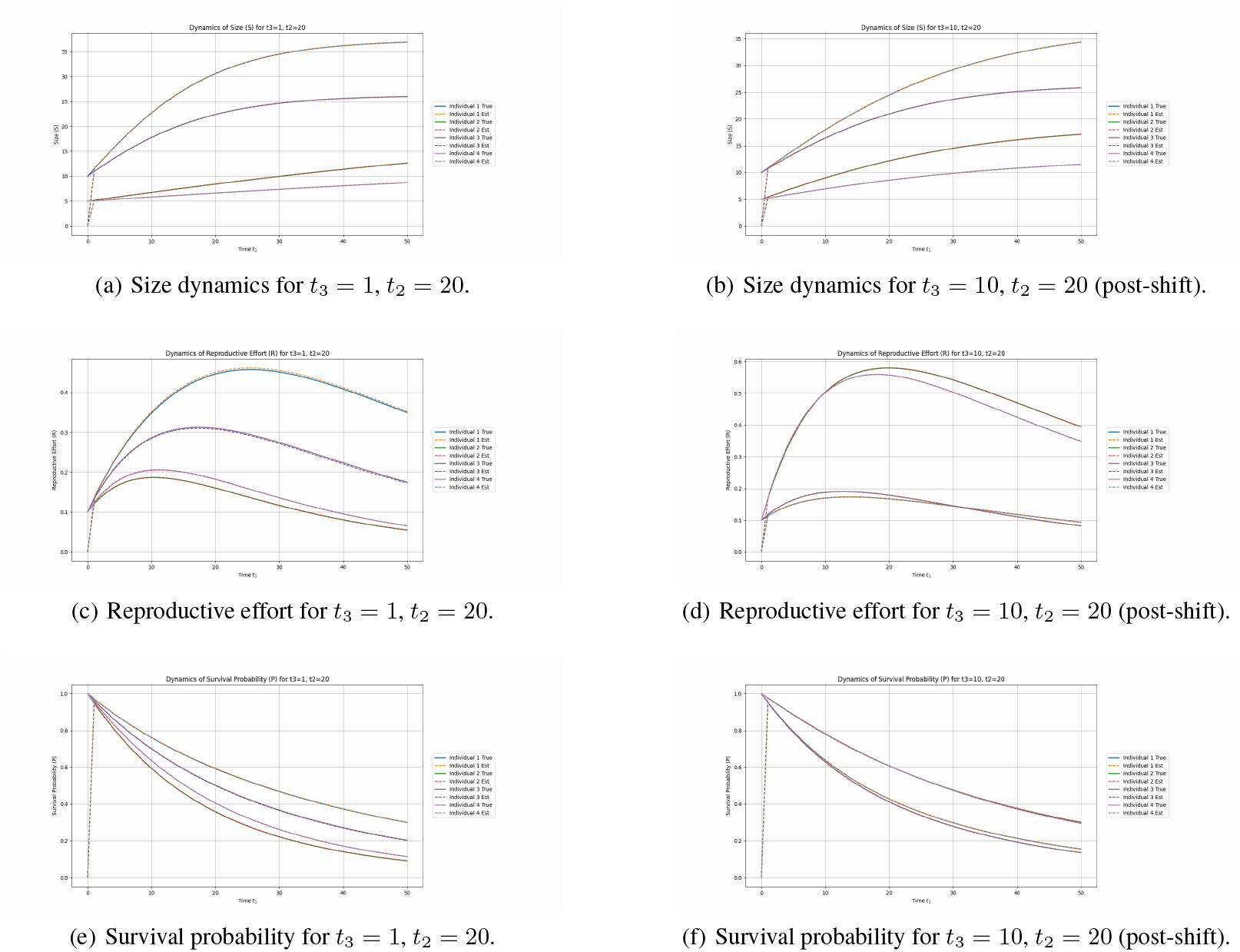
Comparison of size dynamics, reproductive effort, and survival probability for *t*_3_ = 1 and *t*_3_ = 10 at *t*_2_ = 20. The figures highlight the impact of multiscale feedback on the individual life cycle.

Figures 2(c) and 2(d) depict the true versus estimated reproductive effort dynamics for each individual before and after the environmental shift, respectively. Prior to the shift (*t*_3_ = 1), individual *d* = 1 exhibits the highest reproductive effort, driven by favorable hereditary factor values 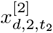 and larger body size 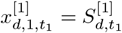. Reproductive effort rises early, peaks around *t*_1_ = 25, and then gradually declines. Individuals *d* = 2, *d* = 3, and *d* = 4 follow similar dynamics but at lower magnitudes, consistent with smaller body sizes and less favorable hereditary factor values. After the environmental shift, individuals *d* = 2 and *d* = 4, who transitioned into more favorable environments, show substantial increases in reproductive effort, especially *d* = 2, who sustains high levels throughout. Conversely, *d* = 1 and *d* = 3, who entered resource-poor environments, exhibit reduced effort relative to earlier trajectories. This demonstrates the continued influence of ecological context in shaping reproductive dynamics. Hereditary factor 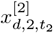 enhances reproductive effort through a multiplicative interaction with size and environmental resource value, but its contribution remains moderate. Notably, these results do not indicate a classical trade-off between reproductive effort and survival probability (see Figures 2(e) and 2(f)). This pattern aligns with a recent meta-analysis in [119] which shows that individuals of higher quality, such as those in better environments or with specific trait values, tend to reproduce more and have higher survival. In the model, reproductive effort and survival reflect a shared developmental and ecological architecture, rather than a strict energetic trade-off. Such patterns are emergent properties of the multiscale feedback structure enabled by the ELC framework. In general, these results align with theoretical work showing that ecological conditions can play a key role in shaping patterns of developmental change [120].

#### 5.2.2 Layer 2: Hereditary Dynamics

The true versus estimated hereditary dynamics for 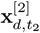 are depicted in Figure 3, highlighting intergenerational changes under varying environmental conditions at *t*_3_ = 1 (pre-environmental shift) and *t*_3_ = 10 (post-environmental shift). Prior to the environmental shift, individuals *d* = 1 and *d* = 3, initially in favorable environments, exhibit the highest values for all three hereditary factors (Figures 3(a), 3(c), and 3(e)). This reflects their elevated 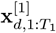 values and greater access to environmental resources. In contrast, individuals *d* = 2 and *d* = 4 exhibit the lowest hereditary factor values due to the harsher environments and lower 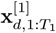 values. Following the switch, we see more pronounced grouping of individuals based on environmental type (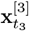 values). In particular, hereditary Factors 2 and 3 exhibit clear divergence between individuals that transitioned into favorable environments (individuals *d* = 2 and *d* = 4) and those that moved into harsher environments (individuals *d* = 1 and *d* = 3) (Figures 3(d) and 3(f)). For Factor 2, values for individuals in favorable environments increase relative to those in harsher environments, reflecting the plasticity of reproductive investment factors in response to improved resource availability. Factor 3, associated with survival factors, shows even stronger environmental grouping, suggesting that survival-related hereditary factors are under tighter environmental regulation or selection pressure. Hereditary Factor 1 (Figures 3(a) and 3(b)) shows a somewhat different pattern. Although individuals initially in favorable environments (individuals *d* = 1 and *d* = 3) start with the highest values at *t*_3_ = 1, after the environmental shift the separation between groups becomes less distinct. Individuals *d* = 2 and *d* = 4, now in favorable environments, show improved trajectories, but the differences across all individuals are more muted compared to Factors 2 and 3. This suggests that Factor 1 may represent a hereditary factor under weaker or slower environmental influence, or one stabilized by internal developmental constraints rather than immediate environmental change.

**Figure 3.**
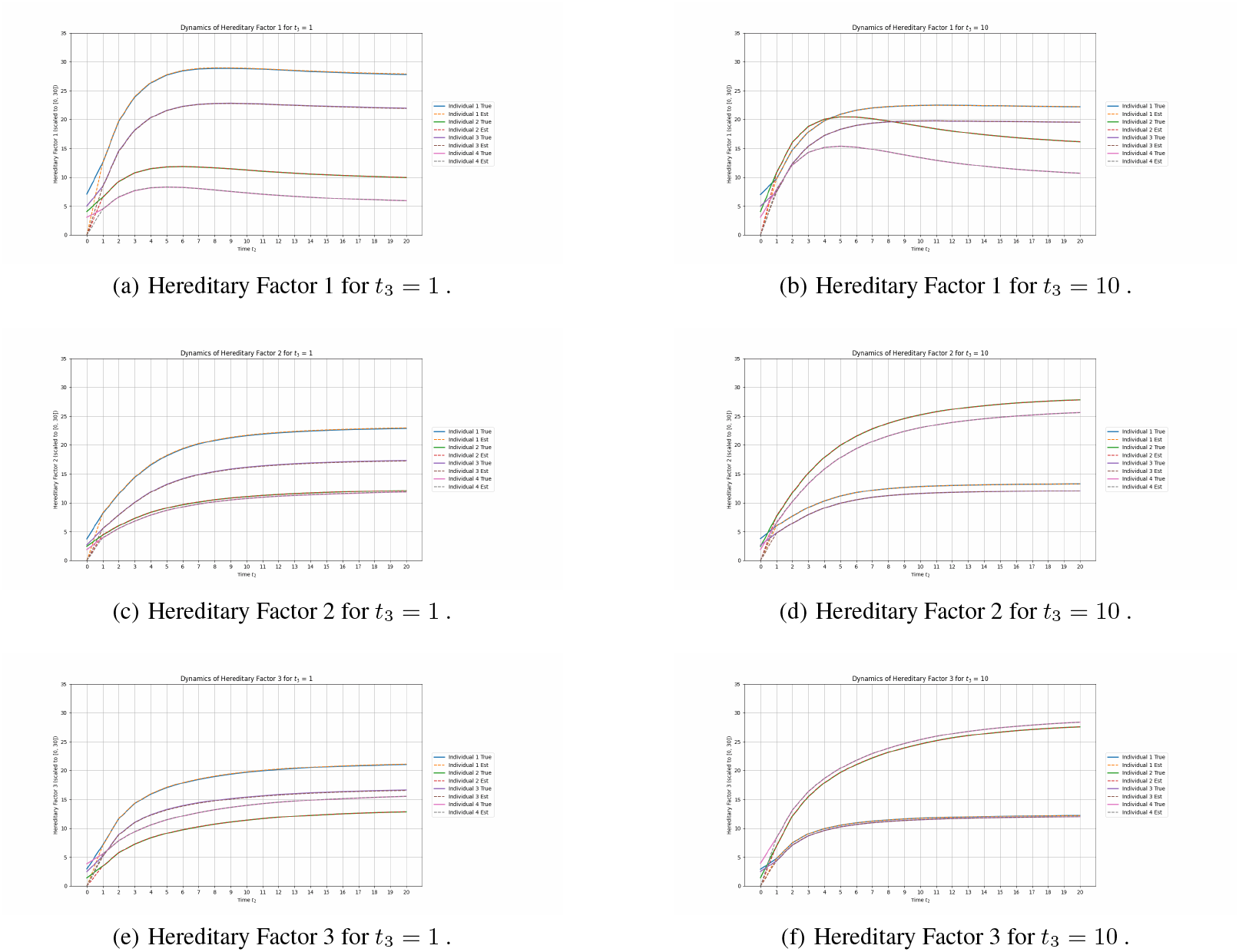
Comparison of hereditary states 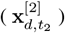 at *t*_3_ = 1 and *t*_3_ = 10. The figures illustrate the impact of environmental shifts on hereditary factors across individuals.

Next, Figure 4 shows the true versus estimated hereditary dynamics with the environmental influence (left) and without the environmental influence (right). When the environmental effect is included (Figures 4(a), 4(c), and 4(e)), individuals cluster more clearly by environmental type, particularly for hereditary Factors 2 and 3. This grouping highlights the strong modulation of reproductive and survival-related traits by environmental conditions. In contrast, when the environmental effect is removed (Figures 4(b), 4(d), and 4(f)), individuals exhibit parallel trajectories, reflecting the influence of developmental dynamics. Specifically, individuals *d* = 1 and *d* = 3 interact through the matrix **B**^[2]^, while individuals *d* = 2 and *d* = 4 interact as well, contributing to the pattern of trait evolution observed without environmental modulation. These patterns highlight the role of developmental dynamics in shaping evolutionary trajectories independently of environmental modulation. Hereditary interactions, encoded through the matrix **B**^[2]^, generate structured correlations among individuals, producing parallel but distinct trajectories even in the absence of environmental effects. The structured divergence in hereditary traits observed under uniform environmental conditions suggests that development itself acts as a source of biased variation, consistent with the idea that developmental processes generate phenotypic variation even in the absence of environmental inputs [121]. As a result, development acts as an intrinsic organizing force, constraining and biasing the pathways of trait evolution and maintaining inter-individual variation across generations.

**Figure 4.**
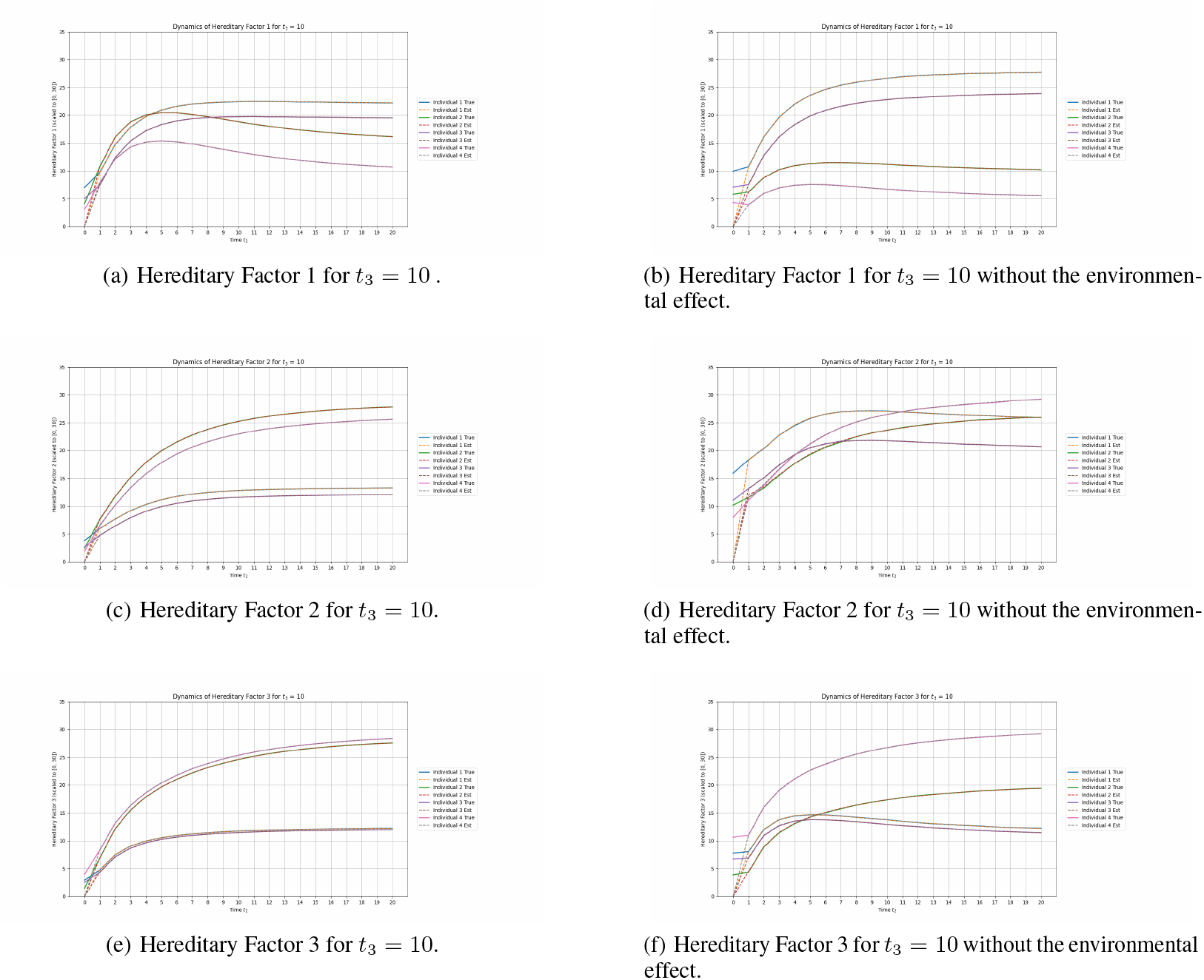
Comparison of hereditary states 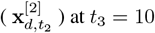 with and without the environmental influence.

Figure 5 compares the true versus estimated hereditary dynamics with developmental influence but no environmental effect (Figures 5(a), 5(c), and 5(e)) against dynamics with neither developmental nor environmental influence (Figures 5(b), 5(d), and 5(f)). When developmental dynamics are included, individuals exhibit coupled divergent trajectories based on pairings encoded by **B**^[2]^ for Factors 1 and 3. A surprising convergence in Hereditary Factor 2 emerges between Individuals 1 and 2, who do not interact and differ in life history and trait optima. This convergence is a geometric outcome of the population-wide structure induced by the interaction topology of **B**^[2]^: its influence propagates through developmental trajectories in a way that aligns certain lineages in state space, even in the absence of direct connections. In contrast, when both developmental and environmental influences are removed, hereditary factor trajectories become highly uniform. Without developmental structure, individuals converge toward similar hereditary factor values, and variation across lineages is reduced. These results demonstrate that developmental dynamics are sufficient to maintain inter-individual variation, acting as an internal organizing force that shapes evolutionary outcomes even in the absence of external environmental inputs. This contrast highlights a core insight of the Extended Life Cycle (ELC) framework: evolutionary outcomes emerge from the interaction between internal developmental pathways and broader hereditary and ecological contexts. Without these nested, reciprocally interacting processes, hereditary factor evolution becomes uniform and uninformative. Thus, this figure illustrates the effects of how the ELC can integrate dynamics across development, heredity, and ecology [122] to produce emergent phenomena.

**Figure 5.**
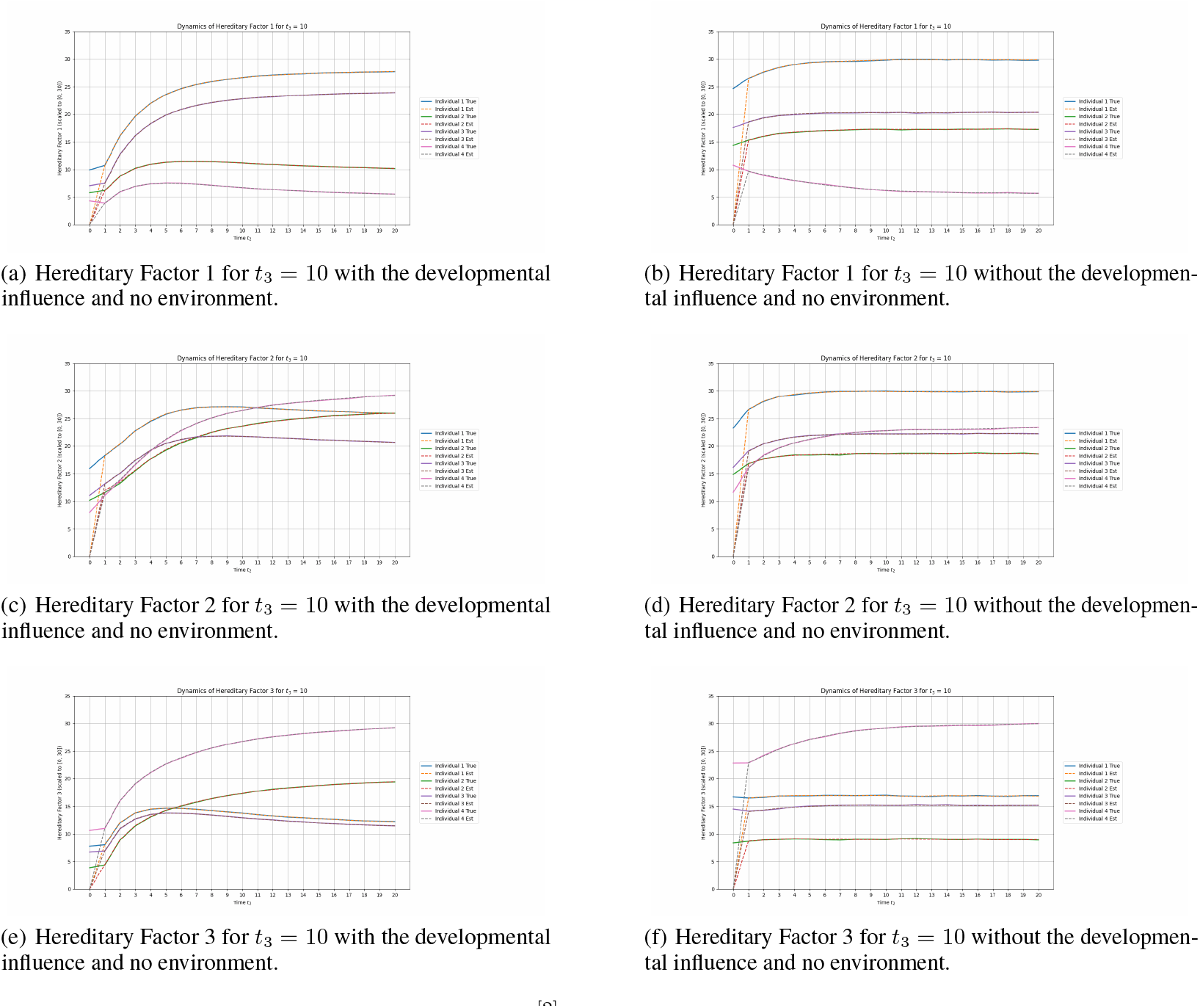
Comparison of hereditary states 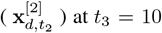 with the developmental influence and no environmental influence (left) and with no developmental or environmental influence (right).

#### 5.2.3 Layer 3: Niche Dynamics

Figure 6(a) shows the true versus estimated environmental state, with a regime shift at *t*_3_ = 5 (black dashed line). For *t*_3_ = 1, …, 4, Components 1 and 2 are favorable, while Component 3 is harsh. Before the shift, Component 1 declines from ≈ 25.0 to 12.0, with a slight recovery driven by Individual 1’s high 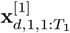 values and corresponding consumption (Figure 6(b)). After the shift, Component 1 transitions to a harsher state and stabilizes at a lower value. Component 2 shows similar dynamics but rebounds more strongly before *t*_3_ = 5 due to Individual 3’s intermediate values. It also declines after the shift, stabilizing around 5.0. Component 3, initially harsh 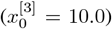, becomes favorable at *t*_3_ = 5, rapidly rising to ≈ 15.5 and stabilizing around 14.0, enabling Individuals 2 and 4 to contribute more. These results demonstrate that environmental dynamics are shaped by life cycle-driven niche construction and external regime shifts.

**Figure 6.**
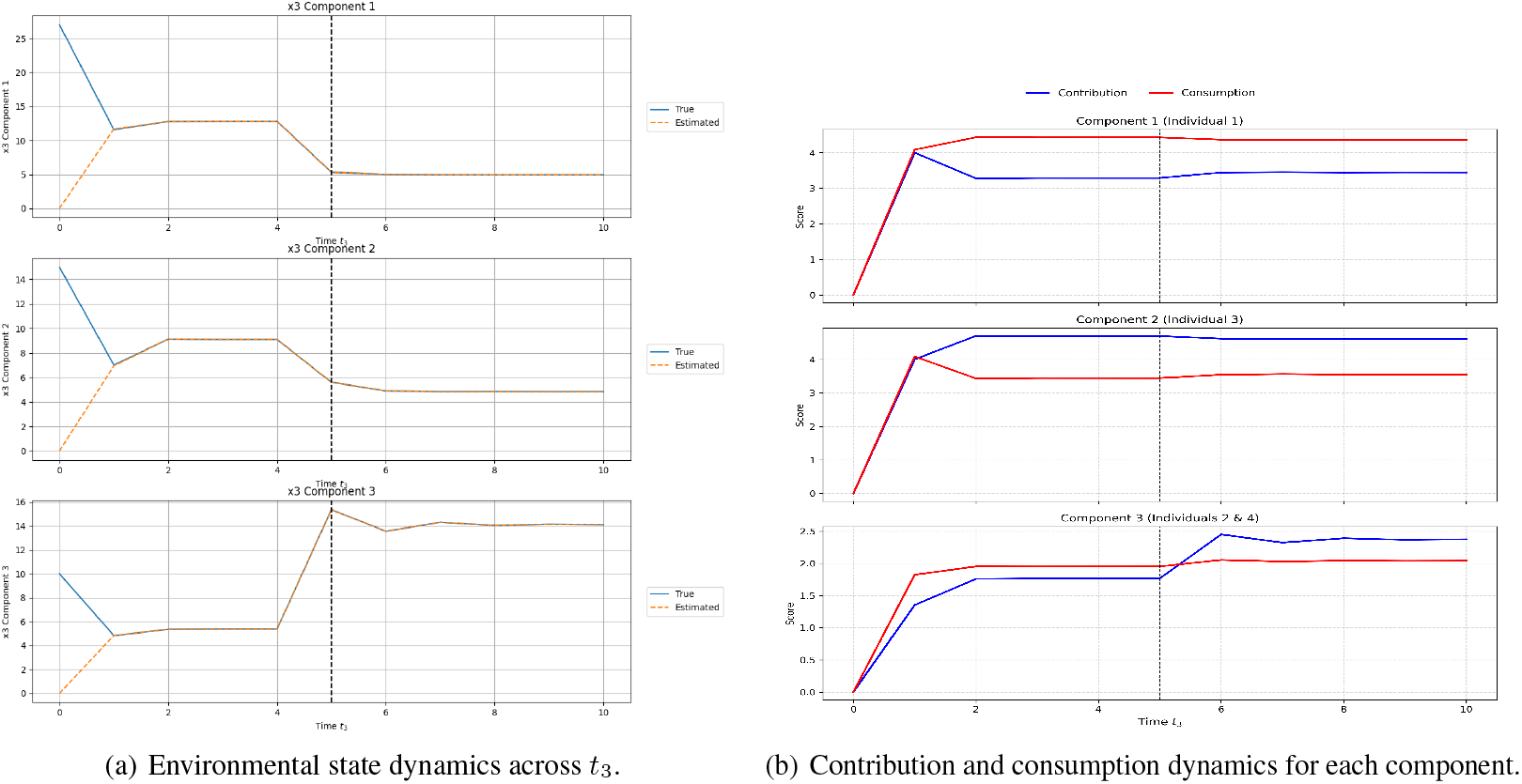
Niche construction dynamics driven by individual life cycle-environment feedback. (a) shows the temporal of environmental components with a regime switch at *t*_3_ = 5 (black dotted line). (b) shows how individual life cycles impact contribution and consumption of environmental resources over time.

Figure 6(b) shows the contribution and consumption dynamics per component. Contribution is modeled as an asymmetric Gaussian centered on the population mean size, with penalties for individuals below the mean; consumption follows a sigmoidal function of size deviation. Individual 2, developmentally closest to the mean, contributes most, with slight decreases after transitioning to harsher conditions. Individual 1, the largest, consumes the most throughout, with a modest reduction post-switch due to a small drop in 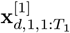. Individuals 2 and 4, being smaller, contribute and consume less overall, but exhibit modest increases when shifted to favorable environments. Individual 3 exhibits the highest contribution due to its optimal 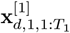 values. These results offer a formal demonstration of niche construction: individuals modify their environments in life cycle-dependent ways, and these modifications feed back into the life cycle dynamics.

## 6 Conclusion

The mosaic of concepts which constitute the EES and EET (non-genetic inheritance, phenotype-led evolution, etc.) is becoming increasingly recognized as a central element of evolutionary theory. However, the success of these emerging theoretical frameworks is dependent on their ability to generate testable theories and predictive models. Historically, the success of the Modern Synthesis was in large part due to the mathematical rigor it gained through the inclusion of population and quantitative genetics. This is the primary challenge that the EES and EET must confront if they are to gain progressive status in the Lakatosian sense [123]. In this work, we have presented one mathematically rigorous approach, termed the Extended Life Cycle (ELC). By formalizing an ontology of causation across multiple time scales, linking phenomena such as individual development, heredity, population structures, and ecological feedback, the ELC moves beyond gene-centrism and unidirectional causation and offers a systems-level view of evolution while incorporating key conceptual elements of the EES and EET. The simulations presented here illustrate how the ELC captures emergent evolutionary dynamics, including life cycle–mediated niche construction, trait divergence and convergence, and trait-environment matching shaped by reciprocal causation. As a modeling platform, it has the potential to be applied across a range of biological contexts and invites further theoretical and empirical work to refine and explore its use.

